# Application of deep neural network reveals novel effects of maternal pre-conception exposure to nicotine on rat pup behavior

**DOI:** 10.1101/2020.07.16.206961

**Authors:** Reza Torabi, Serena Jenkins, Allonna Harker, Ian Q. Whishaw, Robbin Gibb, Artur Luczak

## Abstract

We present a deep neural network for data-driven analyses of infant rat behavior in an open field task. The network was applied to study the effect of maternal nicotine exposure prior to conception on offspring motor development. The neural network outperformed human expert designed animal locomotion measures in distinguishing rat pups born to nicotine exposed dams versus control dams. Notably, the network discovered novel movement alterations in posture, movement initiation and a stereotypy in “warm-up” behavior (the initiation of movement along specific dimensions) that were predictive of nicotine exposure. The results suggest that maternal preconception nicotine exposure delays and alters offspring motor development. In summary, we demonstrated that a deep neural network can automatically assess animal behavior with high accuracy, and that it offers a data-driven approach to investigating pharmacological effects on brain development.

**Significance:** Relating neuronal activity to behavior is crucial to understand brain function. Despite the staggering progress in monitoring brain activity, behavioral analyses still do not differ much from methods developed 30-50 years ago. The reason for that is the difficulty for automated video analyses to detect small differences in complex movements. Here we show that applying deep neuronal networks for automated video analyses can help to solve this problem. More importantly, knowledge extracted from the network allowed to identify subtle changes in multiple behavioral components, which were caused by maternal preconception nicotine exposure in rat pups. Thus, the examples presented here show how neuronal networks can guide the development of more accurate behavioral tests to assess symptoms of neurological disorders.

## Introduction

Animal behavior is a sensitive indicator of brain function. For example, open field exploratory behavioral measures are used to detect changes associated with neurological disorders, pharmacological treatments or environmental conditions (Faraji et al., 2013; Machado et al., 2015; Balbinot et al., 2018; Balkaya et al., 2018; Klahr et al., 2018). Consequently, many methods and tools have been introduced in order to quantify this behavior (Basso et al., 1995; Kabra et al., 2013; Berman et al., 2014; Machado et al., 2015; Wiltschko et al., 2015; Ben-Shaul, 2017; Markowitz et al., 2018; Mathis et al., 2018; Arac et al., 2019; Graving et al., 2019; Pereira et al., 2019). These methods typically require that an experienced observer identifies behavioral abnormalities and designs quantifying measures, which can be time consuming and human designed measures are often sub-optimal due to inherent difficulty of quantifying complex behavior (Schamhardt et al., 1993). To reduce this problem, we introduce a deep neural network that automatically classifies spontaneous behavior and extracts, in a data-driven way, movements that distinguish control and experimental groups of animals.

We applied our network to study the effect of maternal preconception nicotine exposure (MPNE) on rat offspring locomotor development in an open field task (Mychasiuk et al., 2013; Jenkins et al., 2018). Nicotine is a known teratogen capable of perturbing many aspects of development. Nicotine influences brain development by interacting with nicotinic acetylcholine receptors (nAChRs) throughout the brain, affecting cell proliferation, differentiation, and maturation (Dwyer et al., 2008; Blood-Siegfried and Rende, 2010). There is limited research into the effects of maternal nicotine exposure prior to conception (Holloway et al., 2007; Vassoler et al., 2014; Zhu et al., 2014; Yohn et al., 2015; Renaud and Fountain, 2016), and currently no studies consider the impact on early postnatal development. Preconception experience can influence offspring development via three main mechanisms. First, maternal drug exposure may induce physiological changes in the dam that alter the fetal environment, thus resulting in fetal programming even without direct fetal exposure to the drug. Secondly, maternal nicotine exposure may change the quality of maternal care, thereby resulting in the behavioral transmission of an altered developmental trajectory. And thirdly, maternal nicotine exposure may induce epigenetic modifications in the oocyte that shape ontology (Bohacek and Mansuy, 2013).

Because the consequences of MPNE on offspring development are likely to be subtle and vary due to developmental age and the selection of behavioral tests and their measurement, more sensitive behavioral measures could prove useful. For example, the DSM-5 criteria for the diagnosis of ADHD consists of end point measures of the extent to which intrusive activity interrupts behavior in a number of school/home/work settings. Animal models of ADHD similarly use end point measures of open field or swim tests to assess behavioral disruption. We present a new data-driven analysis method to identify MPNE rat pup behavioral changes in an open field task and so provide a better understanding of the neuropharmacological effects of nicotine on brain development. The current research addresses two gaps in our understanding of nicotine’s impact on locomotor activity. First, does nicotine administration during the preconception period in prospective dams, as opposed to the prenatal, preconception + prenatal, or paternal preconception period, affect behavior? Second, is offspring behavior affected at an early stage of development, thus demonstrating an early impact of MPNE on offspring locomotion?

To address those questions, we first analyzed neonatal rat pup video recordings using standard locomotor-derived kinematic measures in order to study an effect that MPNE has on infant rat behavior. Then we showed that a neural network can outperform this conventional analysis in identifying the effect of MPNE. Lastly, we extracted knowledge from the deep neural network in order to identify behavioral components that distinguished the nicotine exposed group from the control group.

This data-driven analysis uncovered novel alterations in “warm-up” behavior (Golani et al., 1981; Eilam and Golani, 1988; Golani, 1992) and provides a new direction in examining the neural consequences of nicotine exposure. This demonstration also highlights a new approach to developing more sensitive behavioral measures for the detection and monitoring of neuropharmacological effects.

## Materials and Methods

### Animals

Procedures were conducted in accordance with the Canadian Council of Animal Care and were approved by the University of Lethbridge Animal Care and Use committee. Animals were given food and water ad libitum and were maintained on a 12-h light/dark schedule (lights on from 07:30 to 19:30) in a temperature- and humidity-controlled (21°/50%) breeding room. A total of 45 female Long Evans born in-house were used. Nicotine-exposed dams (*n* = 23) received 15 mg nicotine hydrogen tartrate salt (Sigma) per liter of drinking water sweetened with 1% sucralose to increase palatability. Control dams (*n* = 22) received 1% sucralose only. Nicotine was administered for seven consecutive weeks beginning in adulthood (90-days-old); seven weeks is the length of the spermatogenic cycle in male Long Evans rats and was chosen to mirror the complementary paternal studies. On average, nicotine-exposed dams consumed 2.4 mg of nicotine per kg of body weight per day across the seven weeks. Females were bred with non-drug-exposed male Long Evans rats (*n* = 45) the day following completion of nicotine administration. Animals in this analysis were pups from 32 successful litters for a total of 351 pups. Eighteen litters (191 pups: 102 female and 89 male) of the animals were from sucralose-exposed dams, and 14 litters (160 pups: 76 female and 84 male) were from nicotine-exposed dams.

### Behavioral testing

Pups were tested in the open field task on post-natal days 10. The testing apparatus was a clear Plexiglas box measuring 20cm × 30cm with a grid of 150 squares (10 squares × 15 squares) on the floor each with a size of 2cm ×2cm (Fig. 1A). Each day, pups were placed individually in the center four squares (shaded black) and left to explore the box for approximately 1 min, while being recorded from above. The open field was cleaned with Virkon between animals.

**Figure 1.**
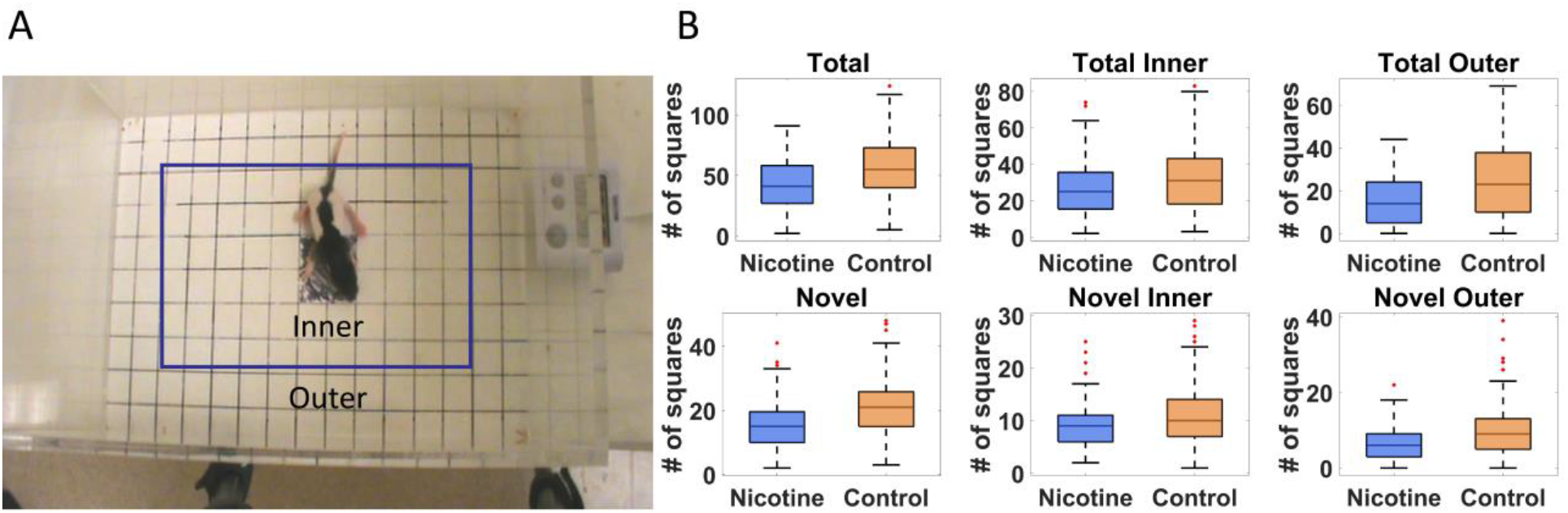
Movement of 10-day old rat pups in open field task. (A) Definition of outer and inner portions of the open field. (B) Comparing movement measures in nicotine and control animals reveals significant effect of MPNE on offspring exploration (see Methods for measures description). On each box, the central mark indicates the median, and the bottom and top edges of the box indicate the 25th and 75th percentiles, respectively. The whiskers extend to ±3SD, and red marks show data points outside of 3SD range.

Kinematic movement measures and their definitions in scoring procedure are as follow: *Novel* = the number of unique squares that either front paw of the pup enters, up to a maximum of 146 (i.e. the box is divided into 150 squares total (10×15), minus the four shaded squares). *Total* = the total number of square entries for either front paw (i.e. the number of times a front paw goes from one square to another)

*Novel Inner* = the number of unique squares in the inner portion of the field that either front paw of the pup enters. (i.e. the number that are within the 6×11 squares in the center of the box, minus the four shaded squares).

*Novel Outer* = the number of unique squares in the outer portion that either front paw of the pup enters. (i.e. the two rows of squares that make up the perimeter of the open field)

*Total Inner* = the total number of square entries in the inner portion for either front paw.

*Total Outer* = the total number of square entries in the outer portion for either front paw.

### Deep Neural Network training and architecture

For training the neural network we used 351 videos: 160 from MPNE animals, and 191 videos from control animals. The original frame rate was 30 frames per second with resolutions 720×480 pixels. However, to reduce the amount of data, we used only every 10th video frame (3 per sec). From each video we excluded the initial period showing the experimenter’s hand releasing the pup, and 50 sec of recording (150 frames) after that was used.

The general network architecture is shown in Fig. 2. First, a pre-trained convolutional neural network (ConvNet) known as Inception-V3 (Szegedy et al., 2016) was used to extract 2048 features from each video frame. This reduced each video to a 2D matrix of the size (2048 features ×150 frames). This matrix was then given as an input to the recurrent neural network (RNN) to predict animal groups. We used a RNN composed of 256 long short-term memory (LSTM) units, which allowed to extract temporal relations between frames. The LSTM layer was followed by a dropout layer of 0.2 to prevent overfitting, and then a dense layer with two neurons with the softmax activation function to classify the animal’s behavior. We used “Group K-Fold” in Keras to split the data randomly and uniformly (to prevent the train and test data being biased) into 5 classes. Each run is initiated with random set of weights. Batch size was 100 and Adam optimizer has been used with binary cross entropy as the loss function. The code for our network including all parameters is available in the Github repository as Behaviour_Recognizer toolbox: https://github.com/rezatorabi13/Behaviour_Recognizer.

**Figure 2.**
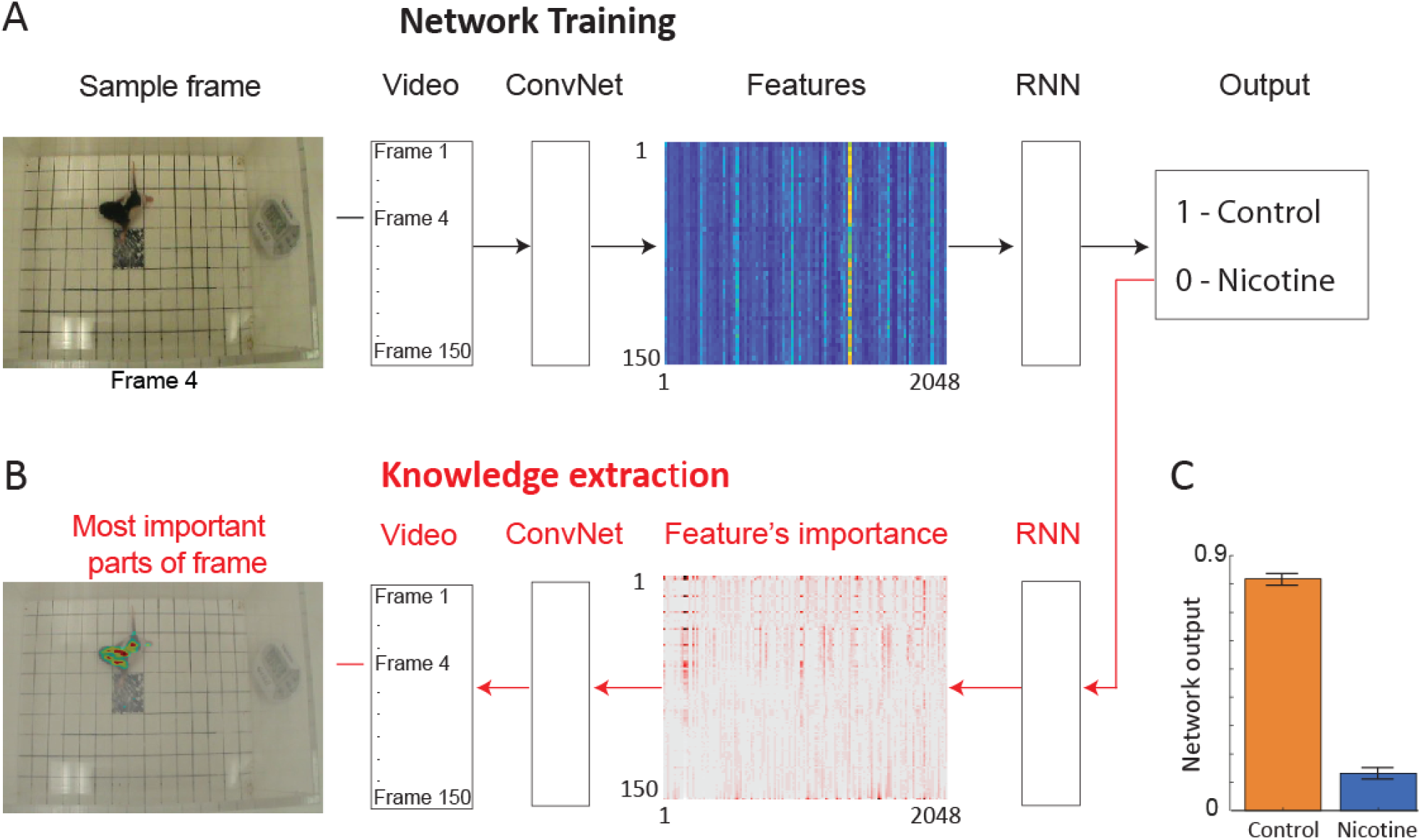
Neural network architecture for data-driven analyses. (A) The network is trained using video clips of single trials (each consisting of 150 video frames). Frames are then passed through a convolutional network (ConvNet) to extract 2048 high level image features from each frame. The features from 150 successive video frames are then given as an input to a recurrent neural network (RNN) to analyze temporal information across frames. Based on this information, RNN predicts a group category for each video clip (Output). (B) After the network is trained, information is extracted from the network weights in order to identify image features and the parts of each video frame that were the most important for network decision making. For visualization, only every 20th feature is shown. (C) Average activity of the output neuron for animal videos from each group.

### Knowledge extraction method

After the network was trained, information was extracted from the network weights in order to identify image features and the parts of each video frame that most contributed to the network decision. For this knowledge extraction from the network, we used the Layer-wise Relevance Propagation method (LPR) (Bach et al., 2015; Lapuschkin et al., 2019) available in the DeepExplain package (Braitenberg and Schüz, 1998). This method uses the strength of synaptic weights and neuronal activity in the previous layer to recursively calculate the contribution (importance) of each neuron to the output score. Because our network is composed of two parts, ConvNet and RNN (Fig. 2A), we first investigated which features were most informative for the RNN to classify animal groups (Fig. 2B middle panel). Next, we propagated feature importance back to pixels in the video through the Inception V3 network (Fig. 2B left panel; Suppl. Fig. 1; Suppl. Text 1). This provided us with information as to which parts of the image the network was ‘attending to’ when making classifications. This allowed to check that the network is indeed using rat movements rather than spurious features, such as the amount of lighting, to discriminate between the treatment groups. Using other methods for knowledge extraction like gradient*input methods (Shrikumar et al., 2017; Ancona et al., 2018) gave qualitatively similar results.

## Results

### Effect of maternal nicotine exposure prior to conception on offspring: analyses of behavior using expert selected measures

Standard locomotor measures were used to investigate the effect of MPNE on offspring locomotor development (Methods). Of 351 rat pups, 191 were from preconception sucralose-exposed dams, and 160 were from preconception nicotine-exposed dams (Methods). When 10-days old (P10), pups were placed singly in the open field for one minute and their behavior was videotaped to investigate locomotor development (Methods). The movement of an animal was described in terms of movement kinematics (Mychasiuk et al., 2013; Jenkins et al., 2018): (1) Total activity - the total number of square entries for either front paw of the animal during the exploration, and (2) Novel activity - the number of unique square entries, which relates to locomotor complexity. Those measures were calculated separately for the inner and outer part of the open field (Fig. 1A, Methods). A statistical comparison of the above movement measures of the MPNE (nicotine exposed dam) and control groups (sucralose-exposed dam) are shown in Fig. 1B. The MPNE group was less active, entered fewer squares and explored fewer novel squares than did the control group (Total_Control_ = 57.0±1.8, Total_Nicotine_ = 42.2±1.6; Total Inner_Control_ = 32.0±1.2, Total Inner_Nicotine_ = 26.4±2.0; Total Outer_Control_ = 24.0±1.2, Total Outer_Nicotine_ = 15.0±1.0; Novel_Control_ = 21.0±0.6, Novel_Nicotine_ = 15.5±0.6; Novel Inner_Control_ = 11.1±0.4, Novel Inner_Nicotine_ = 9.1±0.3; Novel Outer_Control_ = 9.9±0.5, Novel Outer_Nicotine_ = 6.3±0.3; “±” represents SEM; *p*<0.001 for all comparisons using *t*-test; using non-parametric Mann-Whitney U test also gave significant results for all comparisons with *p*<0.003). We did not detect significant sex differences on any of the above measures (t-test, p>0.05 for all measures). In summary, on all measures the MPNE offspring showed less exploration than the control offspring.

### Combining movement measures to distinguish the control vs. nicotine groups

As described above, we quantified the behavior by using typical kinematic measures employed in an open field task. This approach requires assumptions regarding which features of the behavior will be useful in distinguishing between treatment groups. To estimate the reliability of these expert selected features, we used machine learning algorithms to predict treatment group using all six values of behavioral measures described above. We used five different algorithms to ensure that our results are not dependent on a specific data analysis method. For all algorithms we used five-fold cross-validation, where we trained the model on 80% of trials and predicted the treatment group for the remaining 20% of trials. We repeated this process 5 times to predict group category for every trial. The algorithms discriminated between the two groups with accuracy between 57- 64% (Decision tree: 57%; Random forest: 61%; Logistic regression: 61%; K-nearest neighbors: 63%; Support vector machine: 64%) (Suppl. Text 2). This means that based on described movement measures it is possible to tell with about 64% accuracy if it is a control or nicotine group animal (chance level is 50%). We then applied principle component analysis (PCA) to the movement measures. The distribution of points for both classes largely overlapped in PC space (Suppl. Fig. 2). It indicates a weak discriminability between classes, which is consistent with the above result using machine learning algorithms.

### Using deep neural network to distinguish the control vs. nicotine groups

To investigate if additional information could be extracted from the rat pup’s behavior, we used a deep neural network to examine the same videos of MPNE and control animals in the open field task (Fig. 2A). This approach does not require specifying which behavioral measures should be used. Rather, the neural network discovers by itself which features in video (e.g. shapes, movements, etc.) are the most predictive of the treatment groups. Specifically, we used a convolutional network (ConvNet) (Szegedy et al., 2016) to convert each video frame (400×350 pixels) to a set of 2048 features. Those features may loosely correspond to object edges. Features from 150 video frames from a single video clip were then combined and passed to a recurrent neural network (RNN). This allowed an analysis of animal movements throughout each trial (1 trial = 1 video clip consisting of 150 frames corresponding to 50 sec). The network was then trained to assign a correct group category to each video clip (Fig. 2A), and then information was extracted from the network to investigate its decision making (Fig. 2B, see next section).

After training, the network was able to distinguish videos of the MPNE and control groups with 87% accuracy. This accuracy is considerably higher than the classification accuracy obtained from kinematic defined movement features (57%-64%). Fig. 2C shows the average activity of the output neuron for the control group (mean = 0.82±0.02 SEM) and the nicotine group (mean = 0.13±0.017 SEM). The activity of the output neuron was bounded between 0-1, with 1 corresponding to the control category. For example, a value of the output neuron of 0.9 can be interpreted as the network is 90% ‘confident’ that it is a control animal, and only 10% ‘confident’ that it is a MPNE animal. For calculating the network’s prediction accuracy, values of the output neuron above 0.5 was considered as control group, and values below 0.5 as the MPNE group.

To verify that our network does not require fine parameter tuning for robust performance, we also tested four variations of the network. In particular, we modified the number of neurons and layers in the RNN, and we repeated the training and testing on the same data. The modified networks produced results similar to those of the original network (Suppl. Fig. 3). To ensure that network accuracy is not a result of an overfitting and that our network can generalize to new animals, all predictions were obtained using five-fold cross-validation as described above. Thus, no videos of the predicted animal were included in the training dataset. Altogether, these results indicate that there is information about MPNE in the behavior that is not accounted for by the standard movement analyses.

### Extracting knowledge from the neural network

Considering that the network classified the animal groups from videos with higher accuracy than kinematic measures, we investigated what movement features were the most informative for the network. We applied a recently developed Layer-wise Relevance Propagation method (LRP) to extract knowledge from deep neural networks (Bach et al., 2015; Lapuschkin et al., 2019) (Methods). First, we identified which features extracted from the videos were contributing the most to the predictions made by the RNN (Fig. 2B – features importance array), and then we found which parts of each frame corresponded to those most informative features (Fig. 2B – left side). Thus, knowledge extraction reveals the network’s focus for decision making.

Examination of the feature’s importance matrix revealed that certain video frames were particularly informative for the network decision. For example, the first frames had multiple features which contributed to network classification more strongly than subsequent frames (Fig. 3A, Suppl. Fig 4). Plotting the average value of the features separately for each animal group showed the high discriminative power of those initial frames (Fig. 3B). To investigate why, we closely inspected those first frames. We found that on average, there was a meaningful difference in the starting posture and starting movement between the MPNE and control animals. Fig. 3C,D illustrates the difference in their starting posture as soon as the experimenter placed the pups in the open field box. The MPNE animals sprawled, with the fore and hind legs extended, whereas the control animals had their limbs beneath them. In short, the MPNE animals displayed reduced postural support. The lack of postural support indicated by extended limbs could also be observed as the MPNE animals initiated movement. The temporal features of movement were also different between MNPE and control animals as the control animals started to move as soon as they were placed in the open field. They collected their body by bringing their limbs to a weight bearing posture and made small movements of their head as they initiated movement. The MPNE animals mostly lingered (not moving), taking more time to establish postural support and then initiated movement.

**Figure 3.**
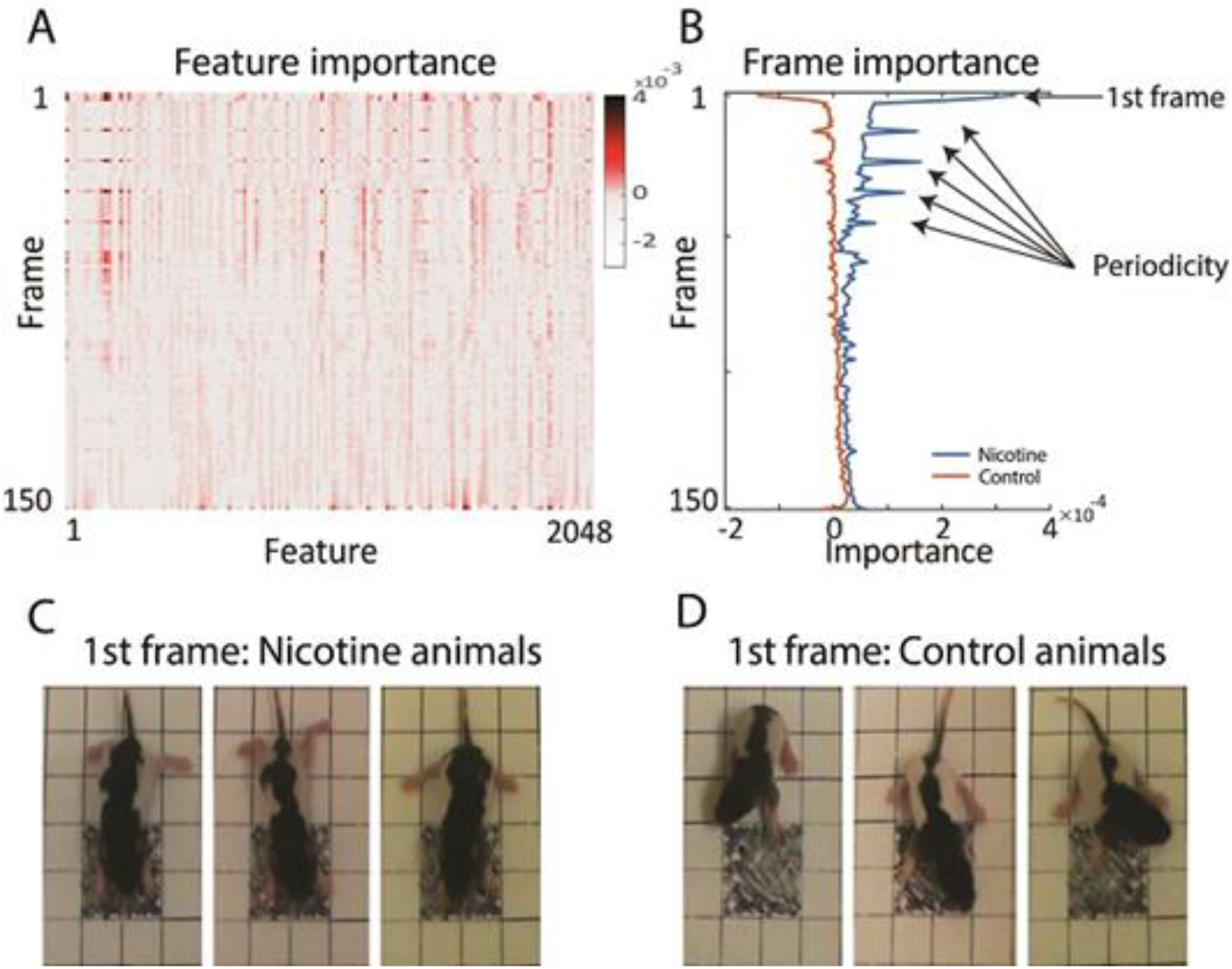
Finding the most informative behavioral features used by the network for decision making. (A) Average feature importance over all videos. (B) Average importance of each video frame, for each animal group. This revealed that the 1st frame and every 11th frame were particularly important. (C & D) Typical starting postures (1st frame) of MPNE (C) and control animals (D). Examples of 3 rats from each group are shown. For visualization clarity, only the portion of frame with the pup is shown. Note extended legs in the MPNE group, and collected legs supporting weight in the control group.

After knowledge extraction from the network revealed that the initial posture is a highly discriminative feature between MPNE and control pups, we developed measure to quantify it. For that, first, we used DeepLabCut software (Mathis et al., 2018), which allowed for semi-automatic marking of position of multiple body parts (four legs, nose, tail base and center of the body; Fig 4A). Next, from *x* and *y* coordinates of resulted marks, we estimated the pose by calculating the average distances between front and hind limbs in the initial frame. Consistent with our visual observation, MPNE animals had a significantly larger distance between front and hind limbs as compared with controls, indicating reduced postural support (Dist_Nicotine_ = 96.2 pixels ±1.53 SEM, Dist_Control_ = 87.57 pixels ±1.46 SEM, p<0.0001, t-test; Fig. 4B).

**Figure 4.**
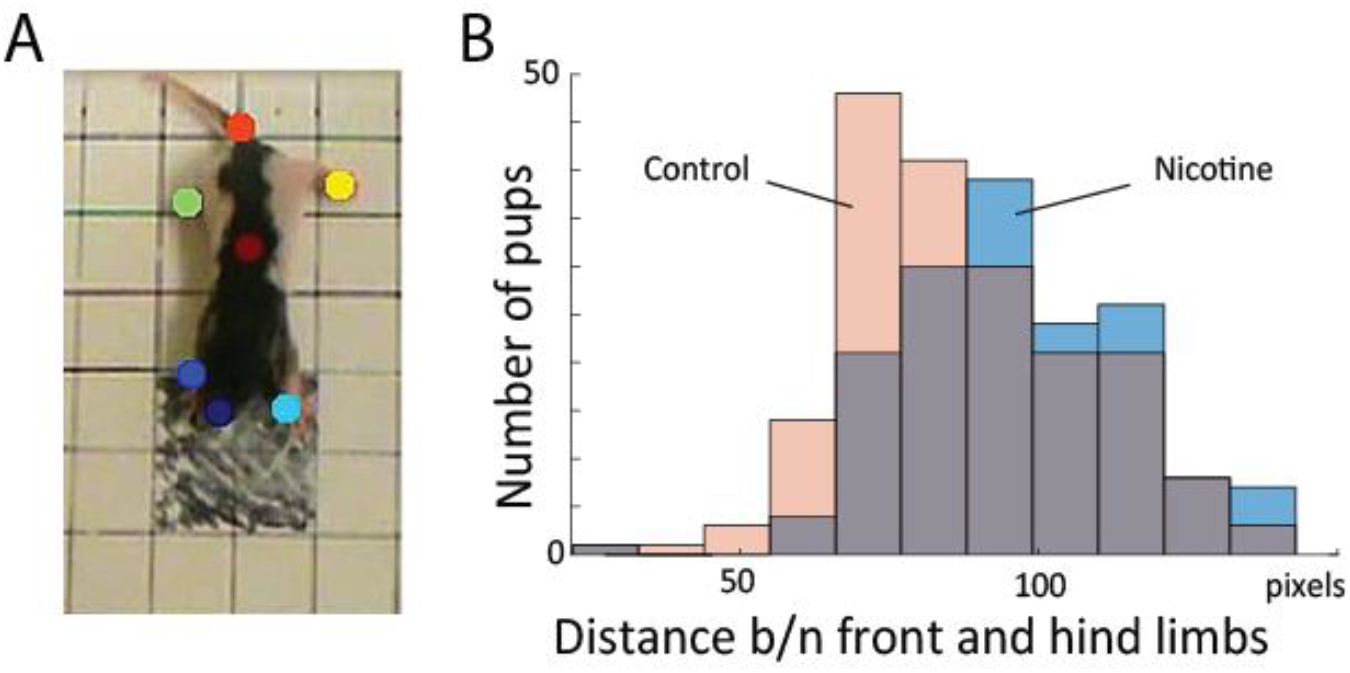
(A) Sample frame with semi-automatically superimposed virtual markers on body parts. (B) Distribution of distance between fore and hind limbs in the first frames for all animals. This shows that MPNE animals (blue bars) had their legs more extended, indicating reduced postural support as comparing to control pups (pink), a result consistent with that shown in Fig. 3C&D.

Visualization of feature importance also showed unexpected periodic changes (Fig. 3B). Specifically, features occurring about every 11th frame, corresponding to period of about 3.7 sec, are particularly informative for the network’s distinction between the MPNE and the control groups. This was also confirmed with spectral analyses shown in Suppl. Fig. 5 & 6. To investigate the behavior underlying the distinguishing movements, we divided the videos into 11 frame segments and aligned the segments (Fig. 5). This revealed a stereotypical, repetitive behavior in MPNE animals. The animals made repeated lateral movements that returned the animal to its initial position. For comparison, Fig. 6 illustrates a typical temporal sequence of two control animals at the same time. The control animals also make lateral movements, but the amplitude and frequency of movement are different from that of the MPNE animals. For example, in the control rat #1, the lateral head movement begins at frame 10 and it ends at frame 18. Its next lateral movement increases in amplitude, thus modifying the sequence of movement (i.e. frames 21 and 31 are not the same in Fig 6 for rat #1). Moreover, some of the control animals pivoted as part of the lateral movement (Fig 6, rat #2). Note that although our analyses showed that features of importance peak at frames 10, 21,…, etc., it should not be interpreted that only those specific frames are of significance to the network. Rather, it should be seen as an indication that at those times the network recognized a specific stereotypical sequence. Thus, the network identified from raw video data the stereotypical behavior as a distinguishing feature of MPNE pups.

**Figure 5:**
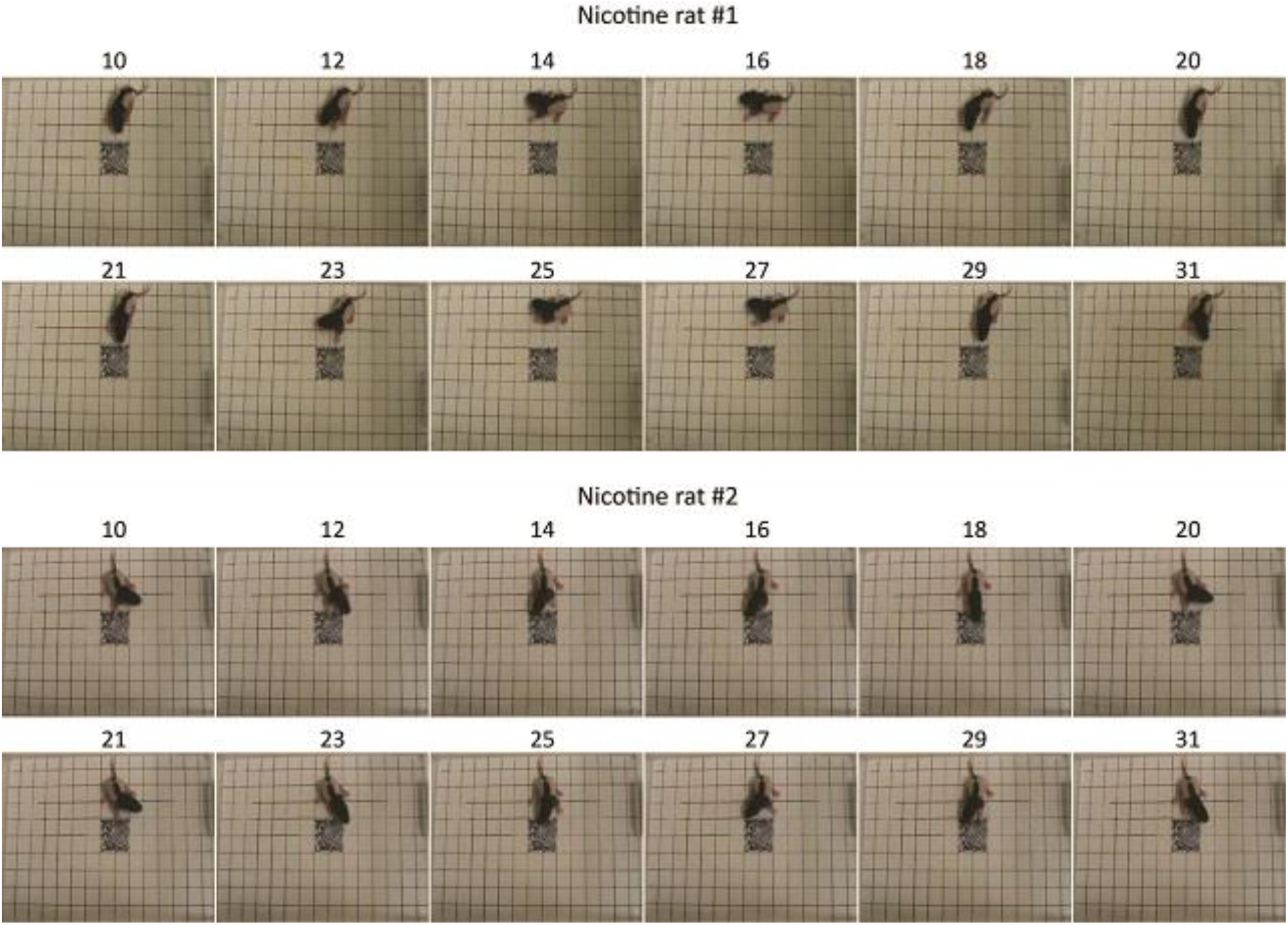
Stereotypical behavior in two rat pups from the MPNE group. Numbers above the pictures represent the frame number. Note the almost exact same body position every 11 frames.

**Figure 6:**
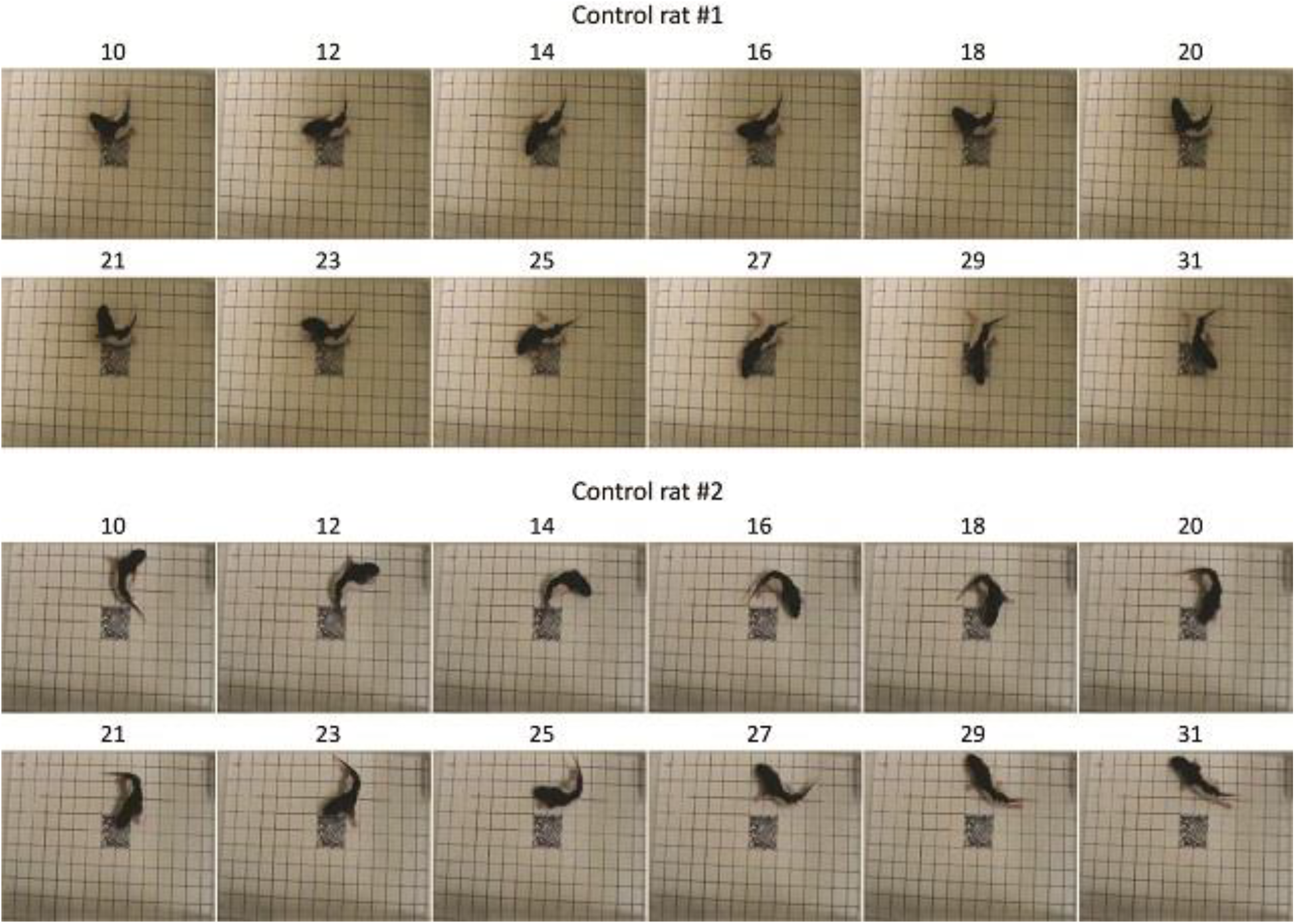
Sample behavior of typical control animals. The movements are less repetitive compared to nicotine animals and more diverse as exemplified by pivoting (rat #2).

To test whether the network discovered differences in stereotypical behavior are significant between groups, we conducted additional analyses. Using DeepLabCut marks (Fig 4A), we tracked nose position for the first 16 frames for all rats (Fig. 7A). Next, we calculated distance between nose position in the 1st and 11th frame. This allowed us to estimate how close to each other was the starting position of the 1st and 2nd sequence of movements. Consistently with results presented in Fig. 5&6, the average distance for MPNE group was significantly smaller (mean Dist_Contr_ = 37.7 pixels ± 2.9 SEM, Dist_Nicotine_ = 60.4 pixels ±3.1 SEM; p<0.0001, Kolmogorov-Smirnov test). Repeating the analyses using the 3rd and 14th frame gave similar results. This confirmed greater stereotypy of movements in nicotine exposed group.

**Figure 7.**
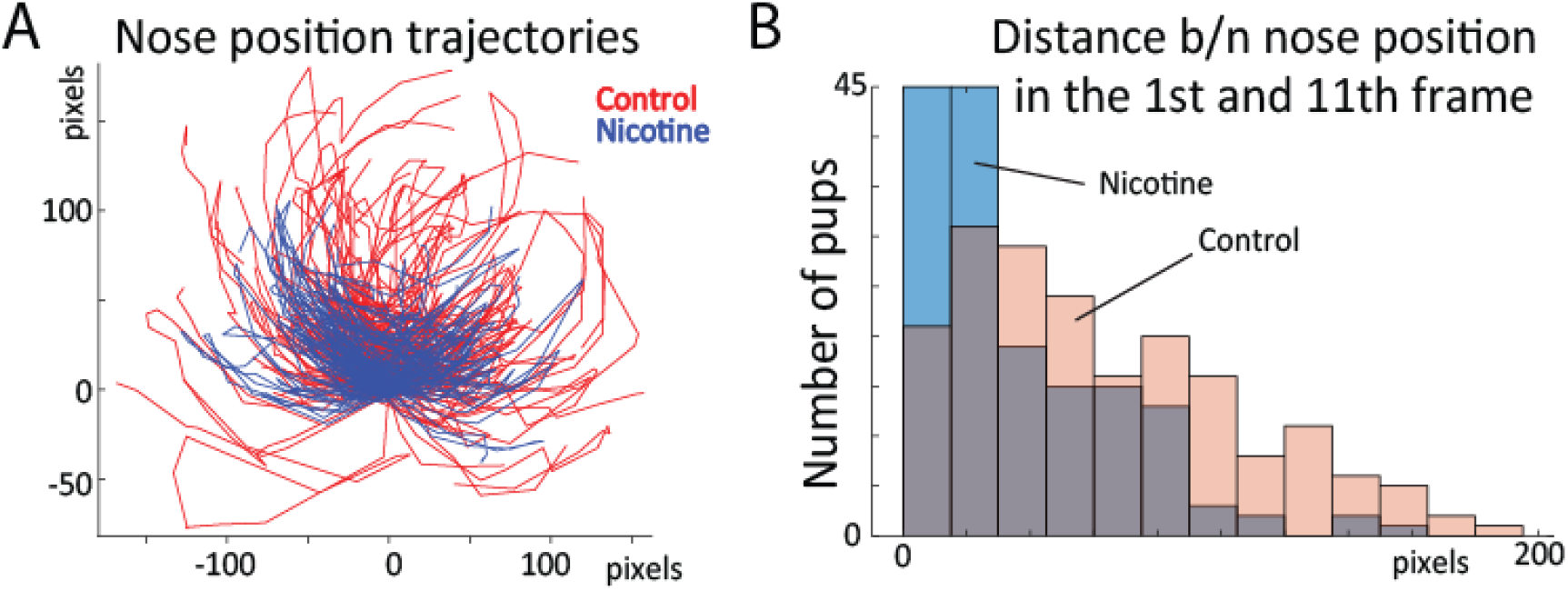
(A) Position of rat nose during first 16 frames for control (red) and MPNE pups (blue). Each animal is represented by a line connecting 16 points. For visualization, only every 2nd animal is shown, and all trajectories are aligned such that nose position in frame 1 is set to point (0,0). (B) Distribution of distances between nose position in frame 1 and 11. The shift of distribution to the left for MPNE group shows that nicotine exposed pups were more likely to return to the same position after first sequence of movements.

We also tested whether only the coordinates of body parts marked with DeepLabCut could provide better features than the ConvNet for predicting the animal group. For that, in all video frames we tracked the position of the nose, limbs, tail base and body center as illustrated in Figure 4A. All points corresponding to frames from one trial were combined as one input to the RNN (similarly to ConvNet features in Fig. 2; RNN with 256 LSTM units). Thus, each video frame was represented by *x*- and *y*-coordinates of seven marked body parts. Using this approach resulted in 62% accuracy to distinguish control and MPNE animals. This is considerably lower accuracy than using original videos (87%), suggesting that ConvNet features selected in a data-driven way, contain additional information useful for behavioral classification. Increasing number of selected body parts may result in improved RNN performance on DeepLabCut features. However, the advantage of our network is that it can directly predict movement deficits from raw videos and does not require human decisions on which body parts to select.

## Discussion

The neural network described here provides a state-of-the-art method for automated behavior analyses. Our network was able to distinguish between control pups and pups whose mothers had consumed nicotine with much better accuracy then standard methods. Presented here a data-driven approach to behavioral analysis offers three advantages. First, it saves researcher time as there is no need for time-consuming human scoring of movement. Second, it increases accuracy, as it explicitly identifies behavioral features that differentiate treatment groups. Third, the largest advantage of the network is that it offers a powerful data-driven approach to gain new insights into behavior. For example, only after looking at feature importance provided by the network did we notice that MPNE pups display altered posture and stereotyped movements that can be diagnosed as early as the first few video frames. Thus, by analyzing the network’s decision-making process, new insights into behavioral differences are obtained. Considering that quantifying behavior is central to most neuroscience research, the method presented here may have a broad applicability.

### Comparison with other methods

One approach to improving the analysis of exploratory behavior is the use of tracking systems using markers that are automatically or manually attached to body parts (Zhou et al., 2008; Parmiani et al., 2019). In the last few years several methods such as LocoMouse (Machado et al., 2015), DeepLabCut (Mathis et al., 2018), JABAA (Kabra et al., 2013), Optimouse (Ben-Shaul, 2017), and LEAP (Pereira et al., 2019) and DLCAnalyzer (Sturman et al., 2020) have been developed to allow users to identify key points in videos, such as the location of a paw, and then it automatically track the movement of those key points across video frames. For instance, in DeepLabCut (Mathis et al., 2018), the experimenter manually labels body parts (e.g. the snout, left ear, right ear, tail) in selected video frames using virtual markers, and then the network is trained to automatically predict the position of those body parts in the rest of the video. Although this method is useful, it requires investigator decisions about relevant body parts, and it requires separate analyses to determine whether the measures are relevant. Here we have also shown that using whole frame video is more informative than using selected markers of body parts to detect pharmacological effects. However, DeepLabCut played a valuable rule in quantifying the behavior allowing us to validate the results we obtained from the knowledge extraction method.

The second category of automated methods such as MoSeq (Wiltschko et al., 2015), MotionMapper (Berman et al., 2014) and (Hsu and Yttri, 2020) first reduce the dimensionality of the video data, and then relate the results to the behavioral components. These methods require image pre-processing and proper image alignment, and additional methods must be applied for classifications. Our method offers an alternative to both of these approaches. First, the convolutional network works with raw images without the need of pre-processing and without the difficult task of image alignment, and second, the network automatically identifies the most relevant behavior for predictions. In short, it offers a one-step solution for feature selection and animal group classification.

Presented here approach also provides significant advancement on our previous network used on the skilled reaching behavior of stroke rats (Ryait et al., 2019), as here we introduced analyses in a time dimension. This allowed us to identify distinctive repetitive movements in rat pup offspring of mothers exposed to preconception nicotine. As movement in time is crucial component of animal behavior, the temporal analyses presented here can help to provide more sensitive measures of neurological deficits.

### Significance of behavioral results

Our data-driven approach identified impaired postural support and increased stereotypical behavior as prominent features of the pups of maternal preconception nicotine exposed (MPNE) mothers. One interpretation of these results is that the MPNE offspring are developmentally delayed compared to the control offspring. One line of research has identified “warm-up” as a key feature of animal movement development. When an animal is placed in a novel location, movement initiation is organized with lateral, forward and dorsoventral movements emerging sequentially and then escalating in amplitude into forward locomotion (Golani et al., 1981; Golani, 1992). Warm-up is also a feature of the ontogeny of motor development in which the topography and amplitude of movement emerges as maturation proceeds (Golani et al., 1981). According to this view, the MPNE animals are motorically underdeveloped relative to the control animals in that their reduced postural support and repetitive movement is diagnostic of a lower level of maturation relative to control pups.

In the future, the behavioural analyses described here could be combined with physiological measures such as electrophysiological recording. Most neuronal analyses rely on using expert selected features of brain activity (e.g.: spike timing, correlations, firing rate in specific time) to relate it to behaviour or sensory stimuli (Luczak et al., 2004; Luczak and Narayanan, 2005; Quiroga and Panzeri, 2009). Applying presented here data-driven approach also to electrophysiological data, may uncover novel features of neuronal activity patterns, more predictive of animal behaviour.

## Acknowledgement

We thank A. Mashhoori, H. Ryait, N. Afrashteh, M. Ello, R. Zabair and J. Karimi for help on different parts of this project.

## Author Contributions

Conceptualization: RT, SJ, IQW, RG, AL; Data acquisition: SJ, AH; Data scoring: SJ, AH; Analyses and interpretation of data: RT, AL, IQW; Writing, Review & Editing: RT, SJ, AH, IQW, RG, AL.

## Abbreviations

MPNE: maternal preconception nicotine exposure

## Supplementary materials

**Suppl. Fig. 1:**
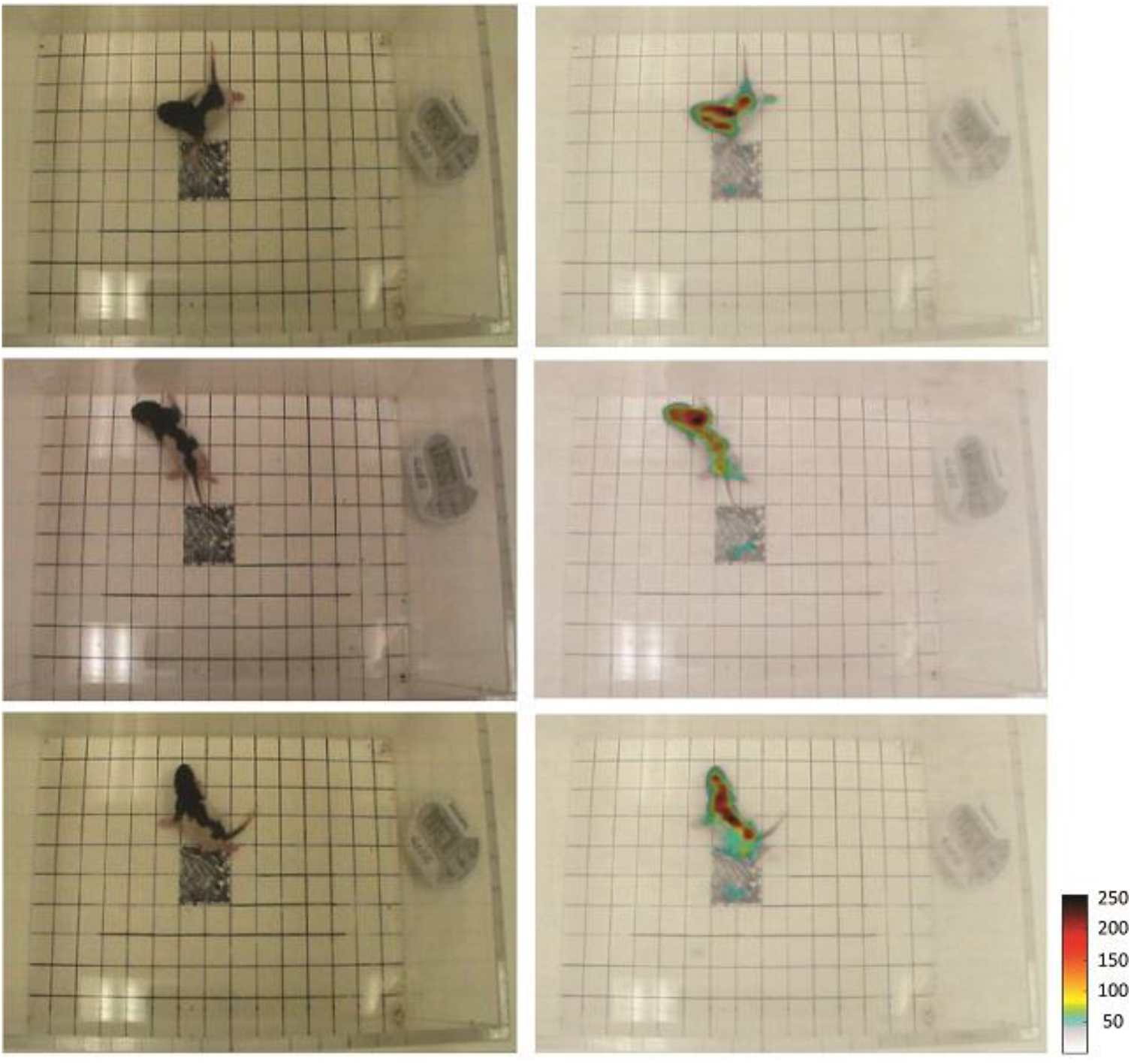
(Left column) Representative video frames, and the same frames with superimposed network focus (Right column). Frames illustrate the most informative pixels used by network to make decisions. This allowed us to verify that the network used features related to rat posture to discriminate control from MPNE animals. This analyses also ensures that the network does not ‘cheat’ by using spurious features like the clock display. To superimpose pixel importance on the frames the values of importance were rescaled to range 50-250. The pixel importance was obtained by using LRP method described above.

**Suppl. Text 1. *Notes on knowledge extraction from neural networks***

Although deep neural networks have demonstrated impressive performance in complex machine learning tasks (Barnickel et al., 2009; Collobert et al., 2011; Le et al., 2011; Ciresan et al., 2012; Montavon et al., 2012; Ji et al., 2013; Szegedy et al., 2015; Krizhevsky et al., 2017) and are key components of many critical decision or predictive processes, they have the disadvantage of acting as a black box, and so not providing information about the internal reasoning for how they reach a certain classification decision (Bach et al., 2015; Lapuschkin et al., 2016; Samek et al., 2017; Srinivasan et al., 2017; Alber et al., 2019). In an attempt to overcome this shortcoming, multiple analysis methods have been proposed to explain predictions of complex non-linear classifiers in terms of input variables (Bach et al., 2015; Samek et al., 2017; Ancona et al., 2018; Alber et al., 2019). One of the most powerful methods in this regard is Layer-wise Relevance Propagation (LRP) (Bach et al., 2015), which we used here. LRP algorithms operate by propagating the prediction backward in the neural network and quantifying the “importance” of input components in classification decisions by attributing relevance scores to them. Video data contain both special (pixels) and temporal (frames) information, so the relevance data provide both special and temporal information about network decision making (Srinivasan et al., 2017; Samek et al., 2018). Special information showed us where the network was looking in the classification task (Srinivasan et al., 2017). This showed that the network is considering the difference in the behavior of control and MPNE animals in classification of video data rather than differences in irrelevant features that may vary between videos (e.g. position of the timer, orientation of the box, or glare from overhead lighting). Temporal information identifies the most relevant time points (Samek et al., 2018) in the behavior. Here, we present a general approach to analyze behavioral video data by identifying the most important spatio-temporal features used by the network for animal classification. This allows us to uncover in a data-driven way the behavioral components most affected by MPNE.

### Suppl. Text 2. *Applying machine learning algorithms on movement measures*

We used five different machine learning methods (logistic regression; decision tree; random forest; k-nearest neighbors; support vector machine) to see how well the movement measures could discriminate between the two groups of animals. In k-nearest method, we tried a different number of nearest neighbors to find the best accuracy. The best accuracy was found for k = 25 (‘k’ is the number of neighbors). In random forest algorithm, the best accuracy was obtained with the number of ensembles n = 50. In SVM, the accuracy with its default parameters is 58%. However, we used grid search for tuning the hyper parameters (C and Gamma) as well as the kernel type to find the best accuracy. The best accuracy was found for C =1, Gamma = 0.01, and polynomial kernel with degree 2.

**Suppl. Fig. 2:**
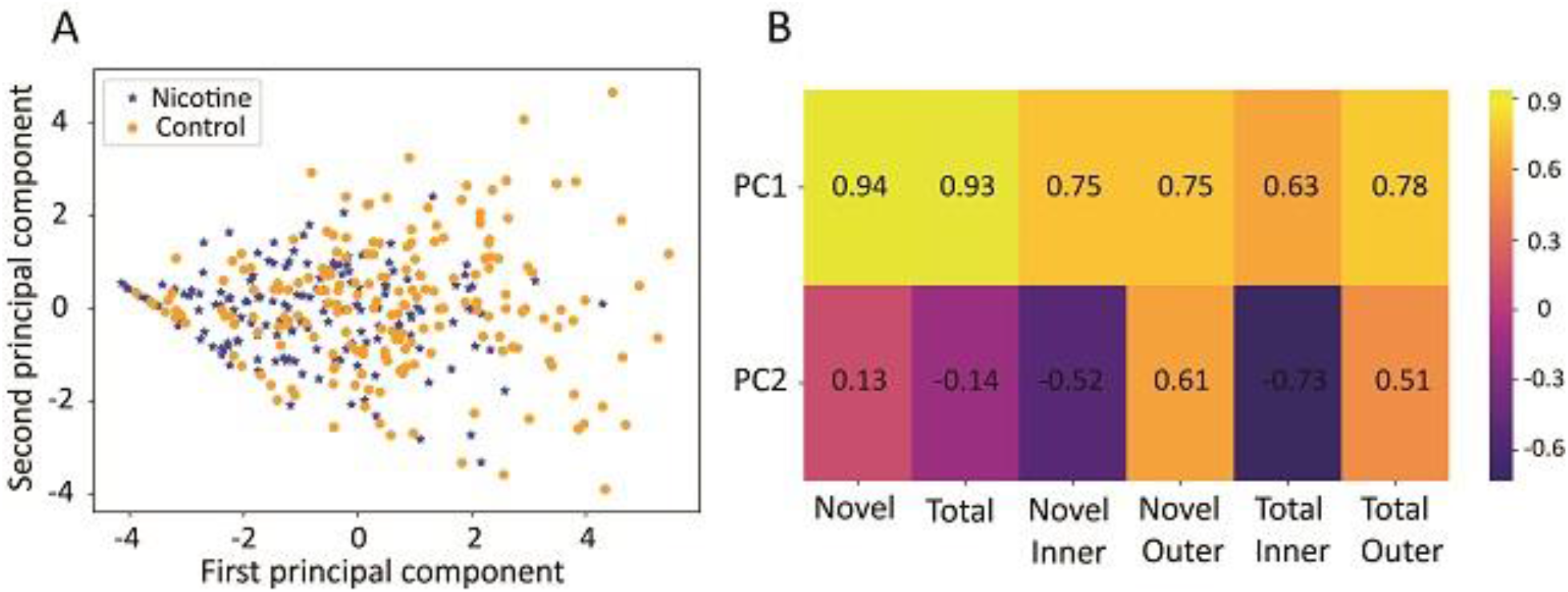
(A) Principal component (PC) space of behavioral measures. Each dot represents a single animal with MPNE group denoted in blue and control group in orange. Note that more control animals have high values of PC1 in comparison with the nicotine animals. To investigate it, we calculated the correlations between principal components and movement measures (B). We found that the first principal component has the largest correlation with the “Novel” and then with “Total” measure. This indicates that control animals have a greater tendency to explore new places (i.e. enter a greater number of unique squares) as well as to explore more (i.e. enter more squares overall) than MPNE animals.

**Suppl. Fig 3.**
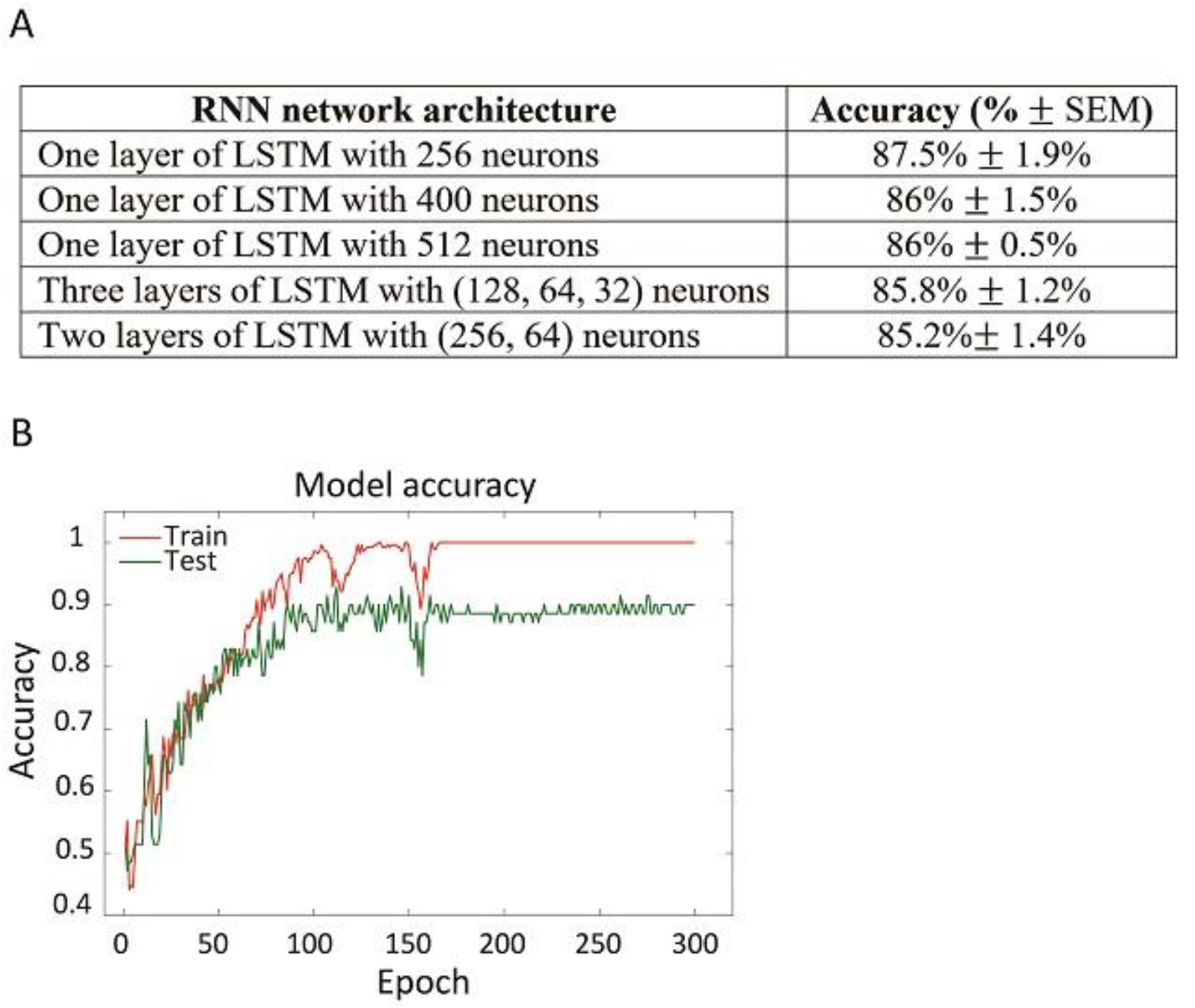
(A) Different architectures of recurrent neural network (RNN) tested. For this manuscript we selected the top network (one layer of LSTM with 256 neurons). However, all tested networks produced similar results, as shown in the table. The network performance was also robust to changes in video preprocessing. Specifically, down-sampling video by taking every 9th frame instead of every 10th frame (Methods) gave similar accuracy of ∼89%. This shows that our network does not need fine tuning to outperform machine learning methods using expert selected movement measures. (B) Sample learning curve from training top RNN with one layer of 256 long short-term memory (LSTM) (Hochreiter and Schmidhuber, 1997; Greff et al., 2017) units. Note that on training data we achieved 100% accuracy (red line), however on testing data accuracy was reduced (green line). This suggests that with larger dataset performance of the network may further improve.

**Suppl. Fig 4.**
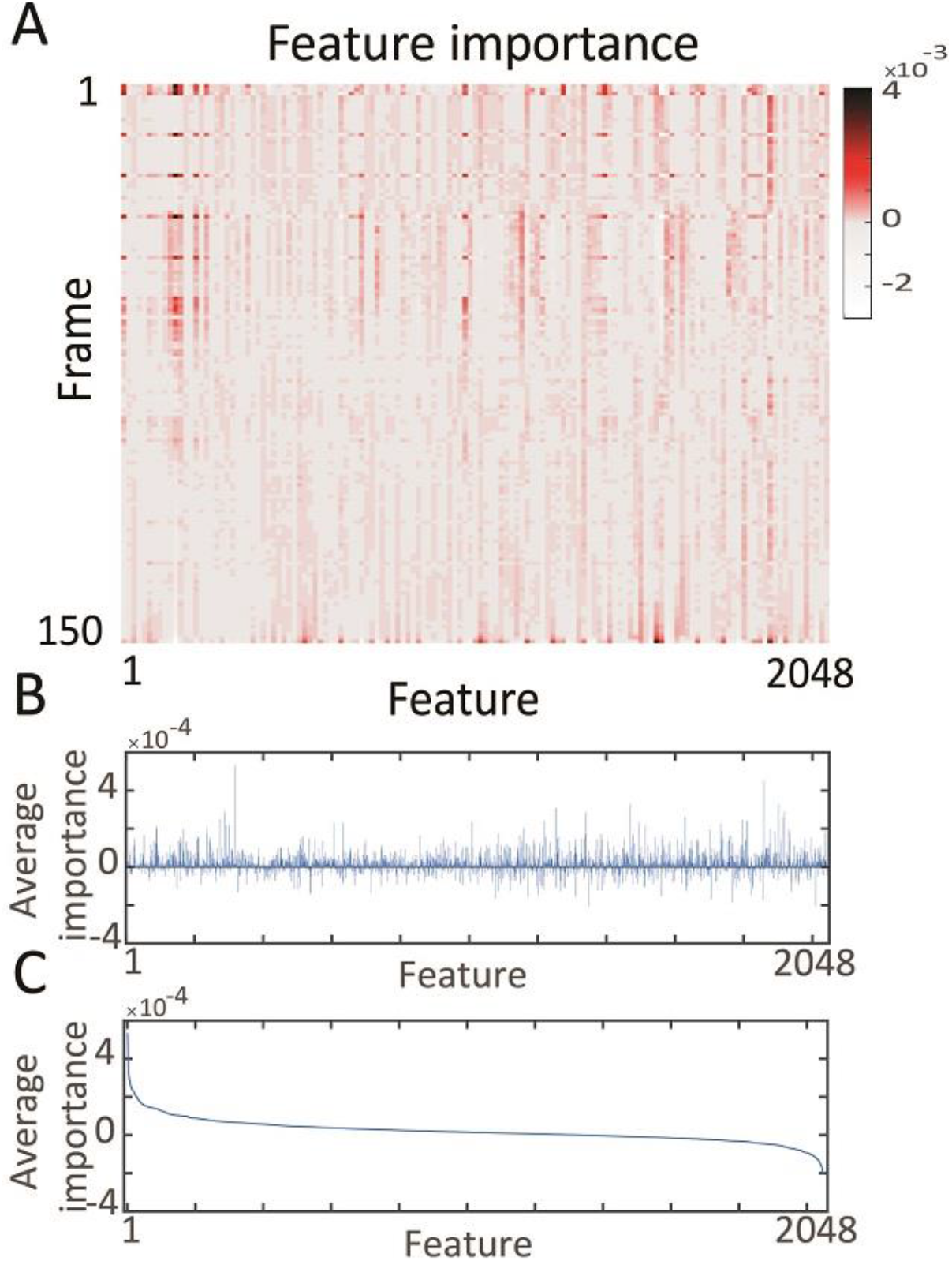
Most informative features for network decisions. (A) Average of feature’s importance over all videos, as shown in main text in Fig 3A. Considering reoccurring peaks in feature importance, it is apparent that the network is identifying periodic behavior, especially in the early 50 frames (∼ 17 sec), to distinguish MPNE animals from control animals. Our RNN network was able to detect a repetitive pattern in the behavior because it is composed of the LSTM units which have memory and are specialized in identifying sequences of activity. (B) Average relevance (importance) of each 2048 features across all video data. Average relevance was obtained by averaging columns in the matrix shown in panel (A). (C) The same average feature importance as in (B), but sorted from highest to the lowest value. It illustrates that about 20 features had a disproportional effect on network decision making.

**Suppl. Fig. 5.**
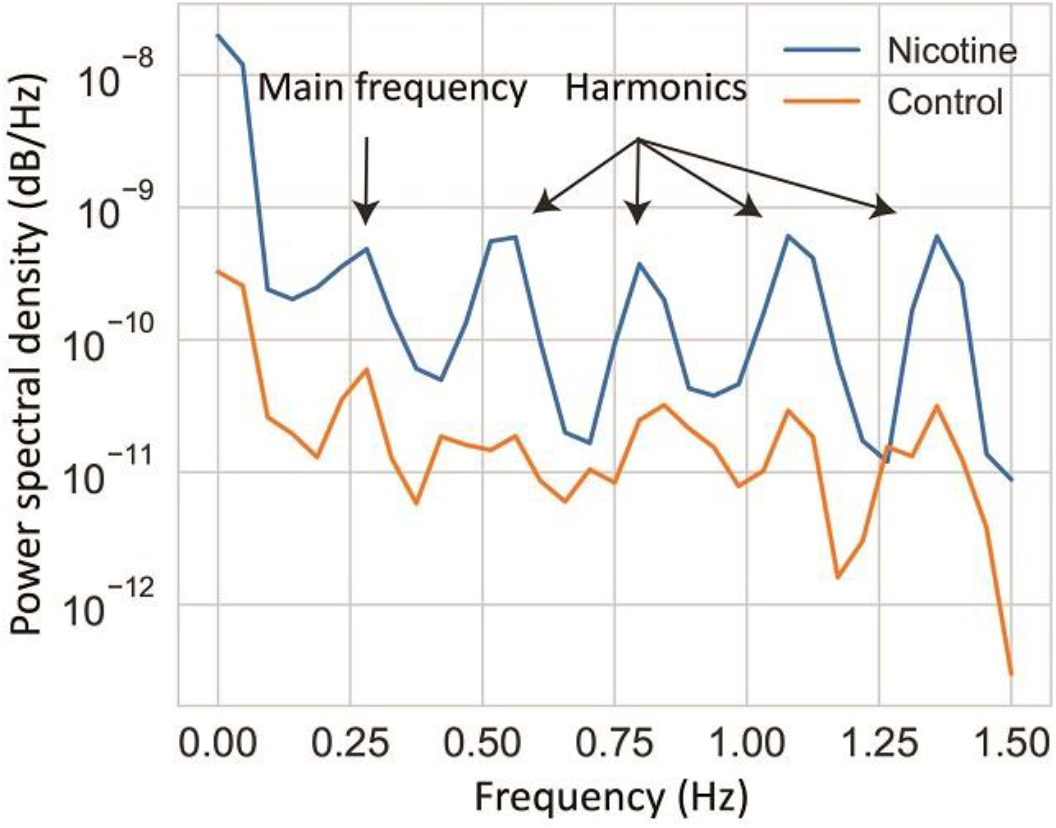
Power spectra analysis. In order to investigate periodic behavior shown in Fig. 3B, we calculated power spectra of the average feature importance. Blue and orange lines denote MPNE and control animals, respectively. Peak in power spectra at frequency of 0.27 Hz confirmed that video features oscillate with period of 1/0.27=3.7s. Note that this periodic behavior was seen mostly in MPNE animals. This indicates that MPNE animals have much more stereotypical behavior, while control animals have more diverse and less repetitive movements.

**Suppl. Fig. 6.**
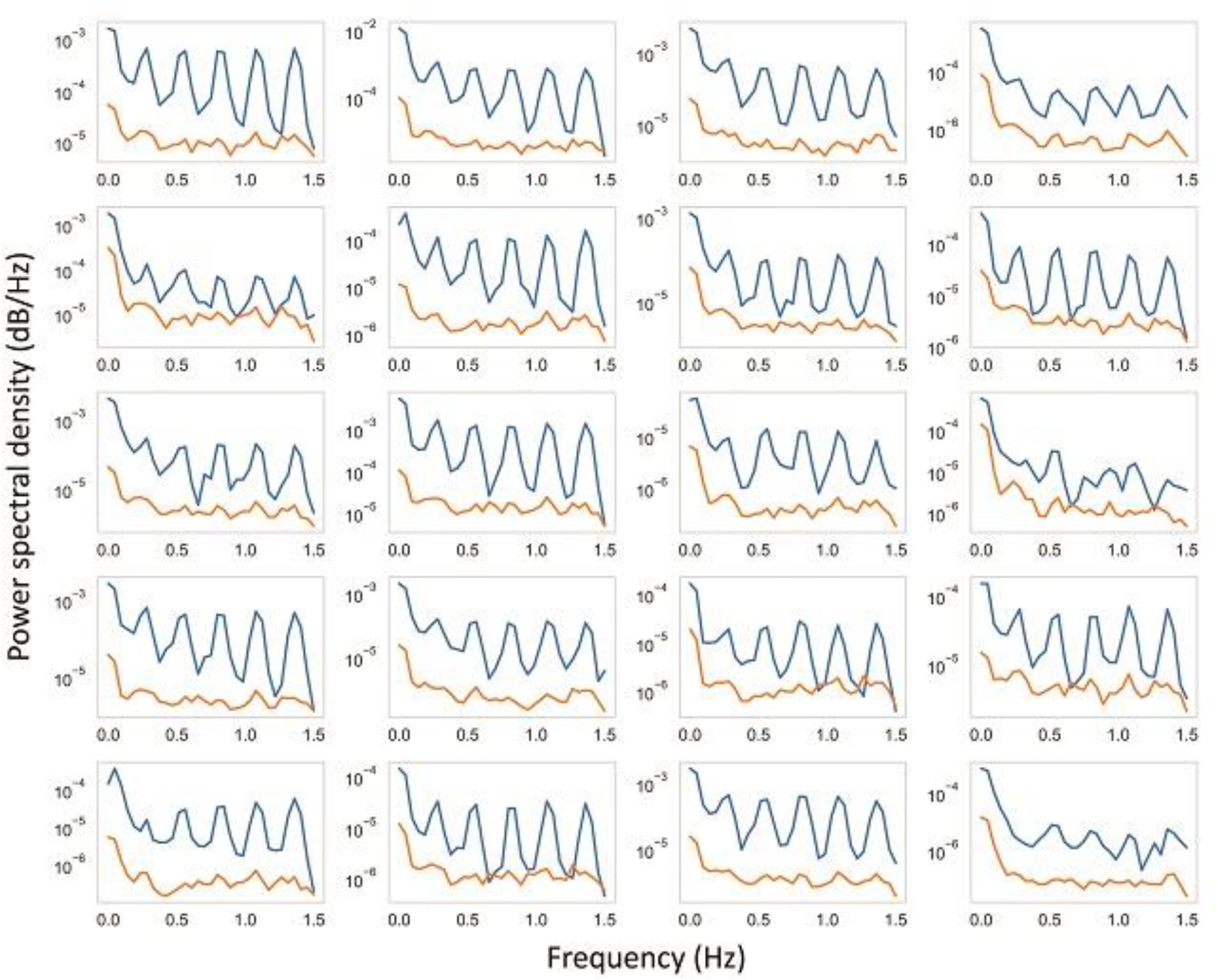
Power spectral analysis for 20 most important features. The blue and orange lines indicate MPNE and control animals, respectively. After identifying the most important features as illustrated in Suppl. Fig 4C, the power spectral for each of the 20 top features was calculated in each video. Then, we averaged spectra of each feature, separately for MPNE and control animals, which lead to the 20 graphs shown above. As can be seen, the periodic behavior with the frequency of about 0.27 Hz is clearly visible in nearly all of the features.

